## Supplementary for "Benchmarking tomographic acquisition schemes for high-resolution structural biology"

| Step | Software | Parameters | |
|---|---|---|---|
| CTF Estimation | CTFFind4 v.4.1.8 | Pixel size | 1.3269 Å |
| | | Acceleration voltage | 300 keV |
| | | Spherical aberration | 2.7 mm |
| | | Amplitude contrast | 0.07 |
| | | Power spectrum size | 512 pixels |
| | | Minimum resolution | 30 Å |
| | | Maximum resolution | 5 Å |
| | | Minimum defocus | 10000 Å |
| | | Maximum defocus | 75000 Å |
| | | Defocus step | 500 Å |
| | | Astigmatism | 100 |
| | | For DS VPP def: | |
| | | Minimum phase-shift | 1.22 rad |
| | | Maximum phase-shift | 2.1 rad |
| | | Phase-shift search step | 0.1 rad |
| Dose-exposure correction | Matlab script from (W. Wan 2017) | | |
| High-peaks removal | eTomo v.4.9.2 (ccderaser) | Peak criterion | 10 |
| | | Difference criterion | 8 |
| | | Maximum radius | 4.2 |
| | | Extra-large difference criterion | 19 |
| Cross-correlation alignment | eTomo (tiltxcorr) | Default parameters | |
| Fiducial model generation | eTomo using “Make seed and track” option | | |
| | Seed model (autofidseed) | Default parameters | |
| | Track beads (beadtrack) | Sobel filter | 1.5 – 3 |
| | | Fill seed model gaps | True |
| | | Local tracking | True |
| | | Local area size | 1000 |
| Alignment transformation computation | eTomo (tiltalign) | Do not sort fiducial into 2 surfaces | True |
| | | Rotation solution type | No rotation |
| | | Magnification solution type | Fixed at 1.0 |
| | | Tilt angle solution type | Fixed |
| | | Distortion solution type | Disabled |
| | | Beam tilt | No |
| Preliminary 8x binned reconstruction | eTomo (tilt) | Logarithm of densities | No |
| | | Radial filtering cutoff | 0.35 |
| | | Radial filtering falloff | 0.035 |
| | | SIRT-like filter | 15 iterations |
| Reconstruction | novaCTF | Correction type | Multiplication |
| | | Astigmatism correction | True |
| | | Slab size | 15 nm |
| Binning | Fourier3D | | |

| | | |
|---|---|---|
| VLPs picking | IMOD v.4.9.2 | |
| Generation of subtomograms initial positions and orientations | Matlab script | |
| Subtomogram Averaging | C++ version of original TOM and AV3 scripts | See the table below |
| FSC | Matlab script implementing phase-randomization as described in (S. Chen 2013) | |

| **SA parameters** | Reference | 8x binned | 4x binned | 2x binned | Unbinned |
|---|---|---|---|---|---|
| Box size (pixels) | 36 | 36 | 72 | 128 | 192 |
| Iterations | 20 | 2 | 3 | 2 | 4 |
| Cone angle opening (degrees) | 12 | 48 | 24 12 8 | 8 4 | 4 2 2 2 |
| Cone angle sampling | 2.9 | 5.8 | 3.9 2.9 1.9 | 1.9 1 | 1 |
| In-plane angle opening (degrees) | 8 | 84 | 48 30 16 | 12 6 | 6 4 4 4 |
| In-plane angle sampling | 1 | 6 | 4 3 2 | 2 1 | 1 |
| Low-pass filter (pixels) | 12 | 12 | 12 | 20 31 | 29 32 32 32 |
| High-pass filter (pixels) | 1 | 1 | 1 | 1 1 | 1 10 10 14 |

**Supplementary table 1: Overview of software and parameters used for processing.**

| | |
|---|---|
| Delay time after tilting by basic increment | 10 s |
| Autofocus offset | 0 µm |
| Autofocus | at least every 3 degrees |
| Repeat record if of field lost is more than | 5% |
| Get tracking image when error in X/Y prediction is more than | 5% |
| Track before autofocusing | Yes |
| Align with preview before getting new track reference | Yes |
| Get new track reference if Record alignment differs by more than | 5% |
| Do autofocus when error in focus prediction exceeds | 0.2 µm |
| Keep beam intensity constant | Yes |
| Limit image shift to | 15 µm |

**Supplementary table 2: Setup for tiltcontroller used to collect the continuous scheme.**

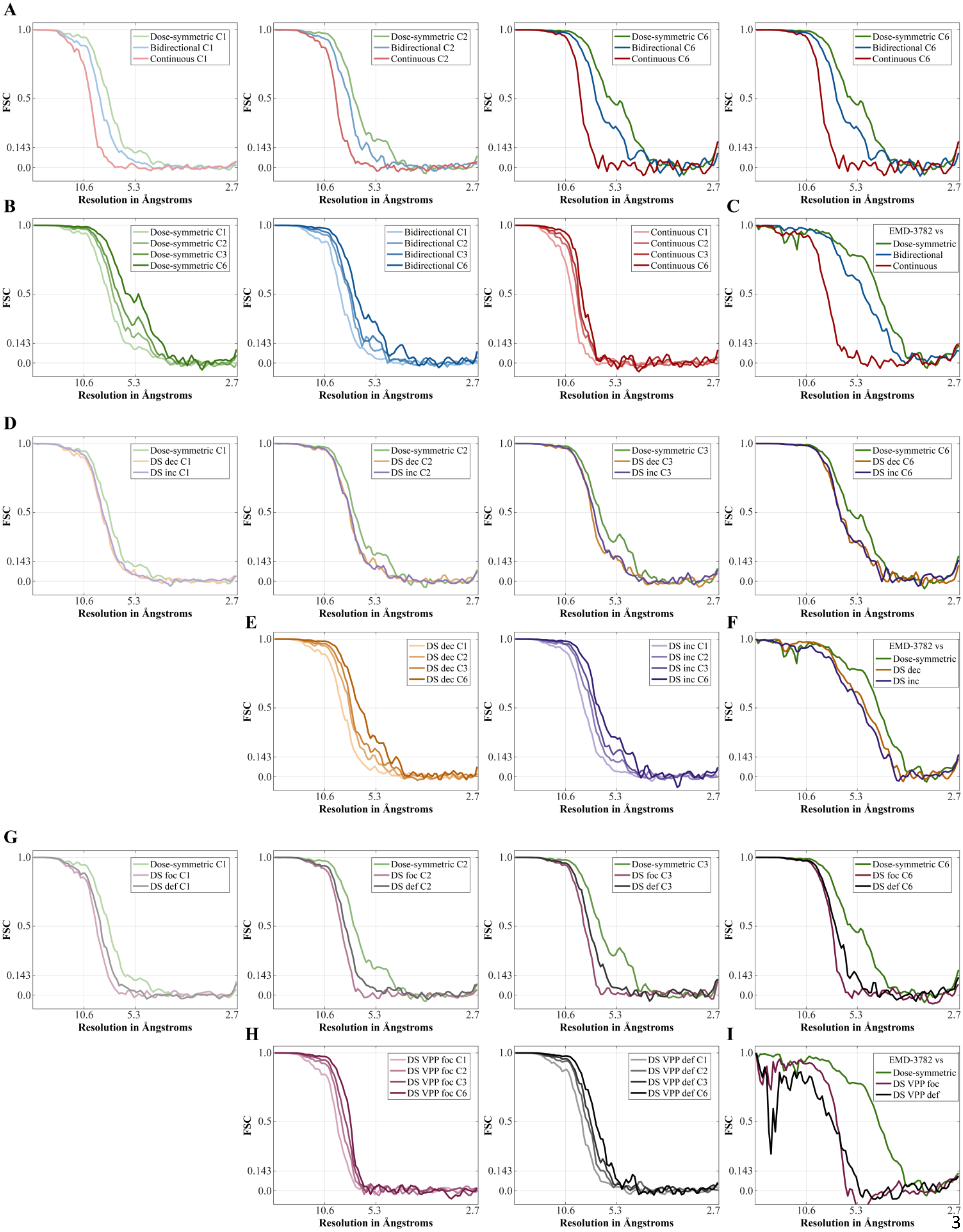
A
Dose-symmetric C1
Bidirectional C1
Continuous C1
Dose-symmetric C2
Bidirectional C2
Continuous C2
Dose-symmetric C6
Bidirectional C6
Continuous C6
B
Dose-symmetric C1
Dose-symmetric C2
Dose-symmetric C3
Dose-symmetric C6
Bidirectional C1
Bidirectional C2
Bidirectional C3
Bidirectional C6
Continuous C1
Continuous C2
Continuous C3
Continuous C6
C
EMD-3782 vs
Dose-symmetric
Bidirectional
Continuous
D
Dose-symmetric C1
DS dec C1
DS inc C1
Dose-symmetric C2
DS dec C2
DS inc C2
Dose-symmetric C3
DS dec C3
DS inc C3
Dose-symmetric C6
DS dec C6
DS inc C6
E
DS dec C1
DS dec C2
DS dec C3
DS dec C6
DS inc C1
DS inc C2
DS inc C3
DS inc C6
F
EMD-3782 vs
Dose-symmetric
DS dec
DS inc
G
Dose-symmetric C1
DS foc C1
DS def C1
Dose-symmetric C2
DS foc C2
DS def C2
Dose-symmetric C3
DS foc C3
DS def C3
Dose-symmetric C6
DS foc C6
DS def C6
H
DS VPP foc C1
DS VPP foc C2
DS VPP foc C3
DS VPP foc C6
DS VPP def C1
DS VPP def C2
DS VPP def C3
DS VPP def C6
I
EMD-3782 vs
Dose-symmetric
DS VPP foc
DS VPP def
FSC
1.0
0.5
0.143
0.0
10.6
5.3
2.7
Resolution in Ångstroms

**Supplementary figure 1. FSC curves. A**. FSC curves comparing the continuous, bidirectional and dose-symmetric schemes with different symmetries. **B**. Comparison of different symmetries for the schemes from A. **C.** Comparison of FSC between C6 maps from the respective schemes and the EMD-3782 map with resolution 3.9Å. **D-F**. Same as A-C, but comparing the dose-symmetric scheme with DS dec and DS inc. **G-I**. Same as A-C, but comparing the dose-symmetric scheme with DS VPP foc and DS VPP dec.

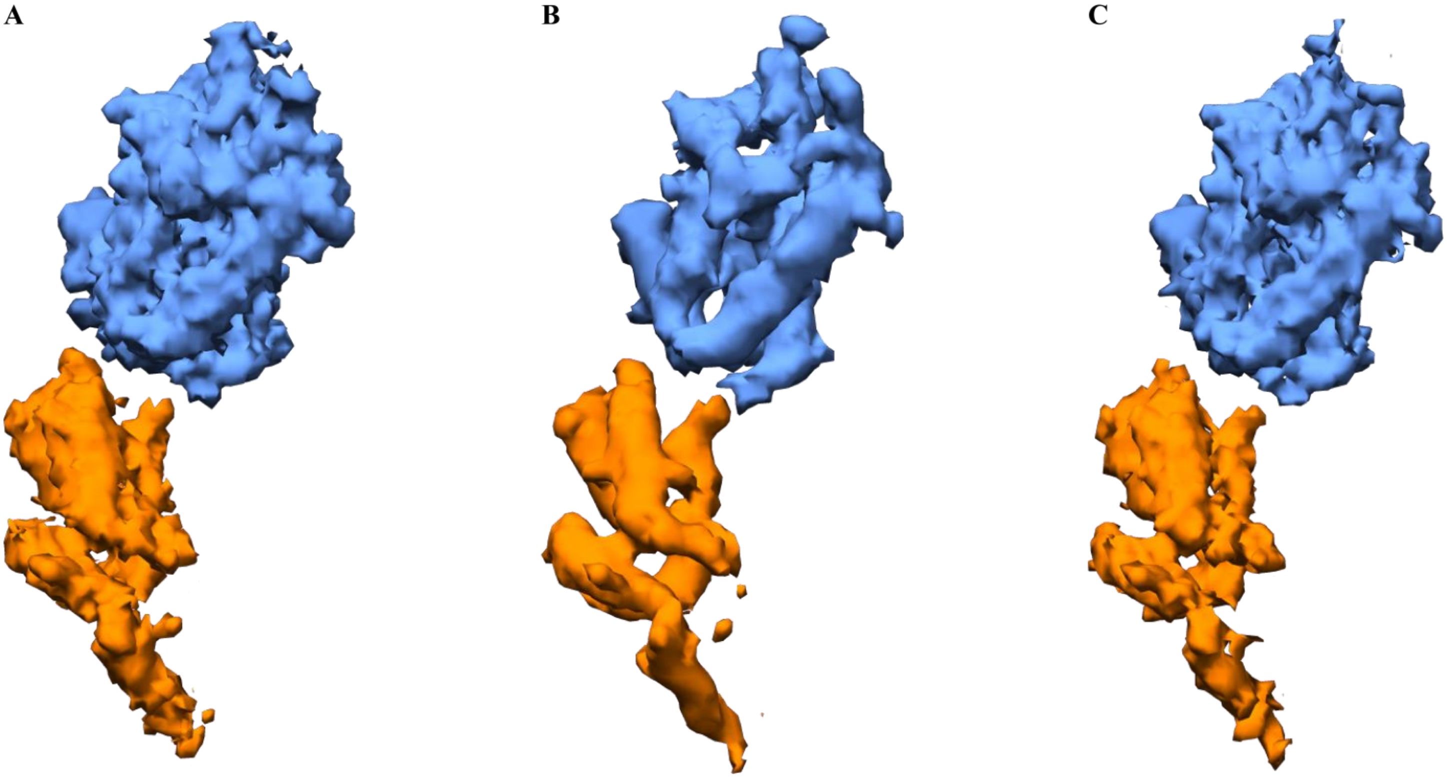


**Supplementary figure 2. Sharpening of VPP structures. A**. A structure of a chain from PDB 5L93 of HIV-1 CA-SP1 monomer obtained by DS VPP foc scheme. **B.** Same as A, but sharpened using empirically determined B-factor of -600. **C.** Same as A, but filtered using the matchto filter from EMAN2 (Tang, et al. 2007). The 4.2 Å structure obtained by the dose-symmetric scheme was used as the reference for the amplitude matching.
